# Estrogen profiling in blood and brain: effects of season and an aggressive interaction in a songbird

**DOI:** 10.1101/2025.06.05.658188

**Authors:** Cecilia Jalabert, Megan Q. Liu, Kiran K. Soma

**Affiliations:** Department of Zoology, University of British Columbia, Vancouver, British Columbia, Canada; Djavad Mowafaghian Centre for Brain Health, University of British Columbia, Vancouver, British Columbia, Canada; Departamento de Neurofisiología Celular y Molecular, Instituto de Investigaciones Biológicas Clemente Estable, Ministerio de Educación y Cultura, Montevideo, Uruguay; Graduate Program in Neuroscience, University of British Columbia, Vancouver, BC, Canada; Integrated Program in Neuroscience, McGill University, Montreal, QC, Canada; Department of Psychology, University of British Columbia, Vancouver, British Columbia, Canada

**Keywords:** catecholestrogens, methoxyestrogens, neurosteroid, estrone, hypothalamus, amygdala

## Abstract

Neuroestrogens are synthesized in the brain and regulate social behavior and cognition. In the song sparrow (*Melospiza melodia*), 17β-estradiol (17β-E_2_) promotes aggression, even during the non-breeding season, when circulating 17β-E_2_ levels are low. Measuring estrogens is challenging due to their low concentrations and the limited sensitivity of many existing assays. Moreover, estrogens other than 17β-E_2_ are often overlooked. Here, we developed a method for simultaneous measurement of eleven estrogens that is highly specific, sensitive, accurate, and precise. We used liquid chromatography tandem mass spectrometry (LC-MS/MS) with derivatization of estrogens by 1,2-dimethylimidazole-5-sulfonyl-chloride (DMIS) to enhance sensitivity. We included four ^13^C-labeled internal standards, which corrected for matrix effects when measuring catecholestrogens and methoxyestrogens. This method greatly improves upon our prior protocol, which could measure only four estrogens and included only one deuterated internal standard. Then, the new method was applied to samples from male song sparrows exposed to either a simulated territorial intrusion (STI) or a control condition during the breeding season or non-breeding season. Only estrone and 17β-E2 were present in blood and brain, while the other nine estrogens in the panel were non-detectable. As expected, there was large regional variation in neuroestrogen levels and very low estrogen levels in blood. There was large seasonal variation, and estrogen levels were lower in the non-breeding season. Despite robust aggressive responses to the STI, estrogen levels did not differ between STI and control subjects in either season. In sum, our novel method enables ultrasensitive measurement of eleven estrogens and will be useful for studies of songbirds and other animals.

## Introduction

Estrogens play critical roles in physiology and behavior. The most widely studied estrogen, 17β-estradiol (17β-E_2_), is traditionally associated with female reproductive behavior, but also modulates memory, attention, executive functions, and multiple social behaviors in females and males [1–7]. In fact, 17β-E_2_ administration rapidly increases aggression in male fish, birds, and mammals [8–12]. Furthermore, social interactions modulate estrogen levels, which may in turn, influence future social behavior [13–16].

The aromatase enzyme (CYP19A1), responsible for estrogen synthesis, is expressed in several brain regions that regulate social behavior [17–21], revealing the capability of local estrogen synthesis. Brain-synthesized estrogens (neuroestrogens) can exert rapid effects near the site of production. Neuroestrogens exhibit rapid fluctuations, as brain aromatase activity is modulated by social interactions, such as courtship [22] and competition [23,24]. Rapid changes in neuroestrogen levels occur even in the absence of changes in circulating estrogen levels. In male zebra finches (*Taeniopygia guttata*), 17β-E_2_ levels in a forebrain region increase rapidly during an interaction with a female, despite stable circulating 17β-E_2_ levels. This effect is abolished by intracerebral administration of an aromatase inhibitor [25].

Song sparrows (*Melospiza melodia*) are a well-established songbird model for studying the steroid regulation of aggression under natural conditions. Males exhibit robust territorial aggression throughout the year, during both the breeding and non-breeding seasons, despite the lack of circulating androgens and estrogens in the non-breeding season [26]. Aromatase is expressed in brain regions implicated in social behavior [27], and increased aromatase activity in the ventromedial telencephalon is associated with elevated aggression in male song sparrows [19]. Additionally, inhibiting estrogen synthesis during the non-breeding season reduces aggressive responses to a simulated territorial intrusion (STI), an effect that is rescued by 17β-E_2_ replacement [28,29]. Furthermore, 17β-E_2_ administration to non-breeding males rapidly increases aggression within 20 min [30].

We recently investigated the rapid effects of an aggressive interaction on brain and blood steroid levels in wild song sparrows [26]. During the breeding season, androgens were detectable but not rapidly modulated in blood or brain regions. In the non-breeding season, androgens were non-detectable in blood of control and STI subjects; however, androstenedione and testosterone increased rapidly in specific brain regions following a STI. Circulating estrogens were not detected during the breeding season, although brain estrogens were present. In the non-breeding season, estrogens were non-detectable in both control and STI subjects. STI did not alter estrogen levels in blood or brain during either season. However, in that study, the assay might not have been sufficiently sensitive to measure very low levels of estrogens in microdissected brain regions. We have very recently developed methods with even greater sensitivity for estrogens.

Liquid chromatography-tandem mass spectrometry (LC-MS/MS) is the gold standard for steroid quantification [31]. Compared to antibody-based assays, LC-MS/MS offers higher specificity and sensitivity and enables the simultaneous quantification of multiple steroids [32–38]. Since estrogens are often present at very low concentrations, assay sensitivity is critical [39,40]. Derivatization can improve ionization efficiency and increase LC-MS/MS assay sensitivity [41–47]. We used an estrogen-specific derivatization reagent 1,2-dimethylimidazole-5-sulfonyl-chloride (DMIS) to quantify four estrogens: estrone (E_1_), 17β-E_2_, 17α-estradiol (17α-E_2_), and estriol (E_3_) in songbird blood, plasma, and brain tissue [45], based on previous studies in humans and mice [44,48]. We also attempted to measure 4-hydroxyestradiol (4OH-E_2_), 2-methoxyestradiol (2Me-E_2_), and 4-methoxyestradiol (4Me-E_2_) but matrix effects were very high for those analytes. In breeding males that were not exposed to an aggressive challenge, E_1_ and 17β-E_2_ levels were much higher in the brain than in the circulation, and local levels of E_1_ and 17β-E_2_ showed regional variation that aligned with aromatase expression. In non-breeding males that were not exposed to an aggressive challenge, we detected 17β-E_2_ only in the caudomedial nidopallium. This study did not include STI subjects. Moreover, this study used only one internal standard (IS), deuterated 17β-E_2_. It would be useful to include additional ISs, additional estrogens, and subjects exposed to an STI.

Here, we improved the estrogen quantification method by using four ^13^C-labeled ISs. We also expanded our estrogen panel by adding four estrogens: 16α-hydroxyestrone (16α-OHE_1_), 2-hydroxyestrone (2OH-E_1_), 2-methoxyestrone (2Me-E_1_), and 4-methoxyestrone (4Me-E_1_), resulting in a panel of 11 estrogens total. Then, using our improved method, we measured estrogens in blood and 10 microdissected brain regions of free-living adult male song sparrows collected during the breeding and non-breeding seasons, with and without an STI.

## Materials and Methods

### Field procedures

The study was conducted on free-living adult male song sparrows during both the breeding season (May 8-24, 2019) and non-breeding season (Nov 2-Dec 2, 2019). Field sites were located near Vancouver, British Columbia, Canada. Subjects were captured using mist nets and conspecific song playback. Immediately after capture, the subjects were deeply anesthetized with isoflurane and rapidly decapitated within 3 min to minimize handling effects. The brain was quickly collected and snap-frozen on powdered dry ice. Trunk blood was collected in heparinized microhematocrit tubes (Fisher Scientific) and kept on wet ice until transported to the laboratory (within 5 h). In the laboratory, both blood and brain samples were stored at ‒ 70°C.

All procedures complied with the Canadian Council on Animal Care, and protocols were approved by the Canadian Wildlife Service and the UBC Animal Care Committee.

### Reagents

High Performance Liquid Chromatography (HPLC)-grade acetone, acetonitrile, hexane, and methanol were from Fisher Chemical. The derivatization reagent, 1,2-dimethylimidazole-5-sulfonyl-chloride (DMIS), was purchased from Apollo Scientific (Stockport United Kingdom, Lot number: AS478881, CAS number: 849351-92-4) [48]. Powdered DMIS was stored at 4°C under nitrogen gas and protected from light and moisture [45]. Once opened, powdered DMIS was aliquoted and stored at 4°C for up to 12 months. On the day of derivatization, acetone was added to individual aliquots of DMIS to prepare a fresh solution of 1 mg/mL. Sodium bicarbonate buffer (50mM, pH 10.5) was prepared in Milli-Q water.

Here, we studied a panel of 11 estrogens: E_1_, 17β-E_2_, 17α-E_2_, E_3_, 4OH-E_2_, 2Me-E_2_, 4Me-E_2_, 16α-OHE_1_, 2OHE_1_, 2Me-E_1_, and 4Me-E_1_. The last 4 analytes were newly added to our previous estrogen panel [45]. All stock solutions were prepared in HPLC-grade methanol.

Certified reference standards for E_1_, 17β-E_2_, and E_3_ were obtained from Cerilliant. 17α-E_2_, 2Me-E_2_, 4Me-E_2_, 4OH-E_2_, 16α-OHE_1_, 2OHE_1_, 2Me-E_1_, and 4Me-E_1_ were obtained from Steraloids.

Since 17α-E_2_ has the same transitions as 17β-E_2_, with retention times differing by only 0.14 min (resulting in overlapping peaks), 17α-E_2_ was included in a separate calibration curve. Calibration curves were prepared in 50% methanol and consisted of 10 points ranging from 0.01 to 20 pg per tube for all analytes, except for 2OHE_1_ and 4OH-E_2_, which ranged from 0.1 to 200 pg per tube. The lower limit of quantification (LLOQ) was determined as the lowest standard on the calibration curve for which the analyte peak had a signal-to-noise ratio >10.

The present study also expanded and improved upon our previous use of IS. Previously, we used only deuterated 17β-E_2_ [45]. Now we employed four ^13^C-labeled ISs, which are more stable and more closely match the retention times of their analytes than deuterated ISs. The ISs used here were 2,3,4-^13^C_3_-estriol (^13^C_3_-E_3_); 13,14,15,16,17,18-^13^C_6_-17β-estradiol (^13^C_6_-E_2_); 13,14,15,16,17,18-^13^C_6_-2-methoxyestradiol (^13^C_6_-2Me-E_2_); and 13,14,15,16,17,18-^13^C_6_-2-hydroxyestradiol (^13^C_6_-2OH-E_2_), all purchased from Cambridge Isotope Laboratories. Each IS was used for analytes with a similar structure and retention time; ^13^C_3_-E_3_ for E_3_ and 16α-OHE_1_; ^13^C_6_-E_2_ for E_1_, 17β-E_2_, and 17α-E_2_; ^13^C_6_-2Me-E_2_ for 2Me-E_1_, 4Me-E_1_, 2Me-E_2_, and 4Me-E_2_; and ^13^C_6_-2OH-E_2_ for 2OHE_1_ and 4OH-E_2_. This grouping ensured optimal correction for matrix effects and variability in recovery across chemically related analytes. Each IS was prepared to a final working solution of 40 pg/mL in 50% methanol.

### Estrogen extraction and derivatization

Estrogens were extracted from brain tissue or blood samples (20 µL) using liquid−liquid extraction as before [26,45,49]. Briefly, 1 mL of acetonitrile was added to each sample, and 50 µL of IS (i.e., 2 pg of each IS) was added to all samples except “double blanks.” Samples were homogenized using a bead mill homogenizer (Omni International Inc., Kennesaw, GA, USA) at 4 m/s for 30 sec. Following homogenization, samples were centrifuged at 16,100 g for 5 min, and 1 mL of the supernatant was transferred to a pre-cleaned borosilicate glass culture tube (12 x 75 mm). Then, 500 µl of hexane was added, and tubes were vortexed and centrifuged at 3200 g for 2 min. The hexane layer was removed and discarded, and extracts were dried at 60°C for 45 min in a vacuum centrifuge. Calibration curves, quality control (QC) samples, blanks, and double blanks were prepared alongside biological samples.

Derivatization was performed as before [45], based on earlier studies [44,48]. Dried extracts were placed in an ice bath and reconstituted with 30 µL of sodium bicarbonate buffer (50 mM, pH 10.5), briefly vortexed, and 20 µL of 1 mg/mL DMIS in acetone was added. The samples were vortexed and centrifuged at 3200 g for 1 min and then transferred to glass LC-MS vial inserts in LC-MS vials (Agilent, Santa Clara, CA, USA). The vials were capped and incubated at 60°C for 15 min, followed by a cooling period of 15 min at 4°C. Finally, the samples were centrifuged at 3200 g for 1 min and then stored at -20°C for a maximum of 24 h before steroid analysis.

### Estrogen analysis by LC-MS/MS

Estrogens were quantified as previously described [45]. Samples were placed in a refrigerated (15°C) autoinjector in a Nexera X2 UHPLC (Shimadzu Corp., Kyoto, Japan) and 35 μL per sample were passed through a KrudKatcher ULTRA HPLC In-Line Filter (Phenomenex, Torrance, CA) and then an Agilent 120 HPH C18 guard column (2.1 mm). Then, estrogens were separated on an Agilent 120 HPH C18 column (2.1 x 50 mm; 2.7 μm; at 40°C) using 0.1 mM ammonium fluoride in MilliQ water as mobile phase A (MPA) and methanol as mobile phase B (MPB). The flow rate was 0.4 mL/min. During loading, MPB was at 10% for 1.6 min, and from 1.6 to 4 min the gradient profile was at 42% MPB, which was ramped up to 60% MPB until 9.4 min. From 9.4 to 11.9 min the gradient was ramped from 60% to 98% MPB. Finally, a column wash was performed from 11.9 to 13.4 min at 98% MPB. The MPB was then returned to starting conditions of 10% MPB for 1.5 min. Total run time was 14.9 min. The needle was rinsed externally with 100% isopropanol before and after each sample injection.

We used two multiple reaction monitoring (MRM) transitions for each analyte and one MRM transition for each IS (Table 1). Steroid concentrations were acquired on a Sciex 6500 Qtrap triple quadrupole tandem mass spectrometer (Sciex LLC, Framingham, MA) in positive electrospray ionization mode for derivatized estrogens and negative electrospray ionization mode for underivatized estrogens. We monitored underivatized estrogens and confirmed that the derivatization reaction was complete. No peaks were detected in the blanks and double blanks.

**Table 1.** Scheduled multiple reaction monitoring for derivatized estrogens.

| Analyte | Retention time (min) | Quantifier m/z | Qualifier m/z |
| --- | --- | --- | --- |
| **E<sub>1</sub>** | 10.46 | 429→365 | 429→96 |
| **17β-E<sub>2</sub>** | 10.74 | 431→367 | 431→96 |
| **17α-E<sub>2</sub>** | 10.88 | 431→367 | 431→96 |
| **<sup>13</sup>C<sub>6</sub>-E<sub>2</sub>** | 10.74 | 437→373 | - |
| **16α-OHE<sub>1</sub>** | 8.25 | 445→381 | 445→96 |
| **E<sub>3</sub>** | 7.97 | 447→383 | 447→96 |
| **<sup>13</sup>C<sub>3</sub>-E<sub>3</sub>** | 7.97 | 450→386 | - |
| **2Me-E<sub>1</sub>** | 10.74 | 459→300 | 459→299 |
| **4Me-E<sub>1</sub>** | 10.42 | 459→299 | 459→97 |
| **2Me-E<sub>2</sub>** | 11.16 | 461→302 | 461→283 |
| **4Me-E<sub>2</sub>** | 10.57 | 461→283 | 461→161 |
| **<sup>13</sup>C<sub>6</sub>-2Me-E<sub>2</sub>** | 10.99 | 467→308 | - |
| **2OH-E<sub>1</sub>** | 10.23 | 603→380 | 603→96 |
| **4OH-E<sub>2</sub>** | 10.62 | 605→382 | 605→96 |
| **<sup>13</sup>C<sub>6</sub>-2OH-E<sub>2</sub>** | 10.62 | 611→388 | - |

Note. For all <sup>13</sup>C-labeled ISSs, only one transition was monitored.

### Assay accuracy and precision

Accuracy and precision were assessed using in-house prepared QCs with known amounts of standard in neat solution at 2 concentrations. The low concentration QC had 0.5 pg, and the high concentration QC had 2 pg of each estrogen, except 2OHE_1_ and 4OH-E_2_ which had 5 and 20 pg respectively. For 17α-E_2_ only the low 0.5 pg QC was used. Biological samples were analyzed across six different assays, with three replicates of each concentration in each assay (n=36 QCs total). Accuracy was determined by comparing the measured values to the known values. Precision was determined by calculating the coefficient of variation of QC measurements within the same run (intra-assay variation) and across different runs (inter-assay variation).

### Matrix effects

Matrix effects were assessed in blood samples (20 µL) and brain samples (1.5 mg) from a total of n=5 animals. First, we evaluated estrogen recovery from biological matrices. To this end, pooled blood and pooled brain samples were each divided into two aliquots: one (unspiked) was analyzed to determine endogenous estrogen concentrations, and the other was spiked with a known amount of standard prior to extraction to determine the effects of each biological matrix on steroid recovery. A third tube, containing the same amount of standard in a neat solution (i.e., 50% methanol solvent only), served as the reference. Each tube was assessed in 5 replicates.

Recovery (%) was calculated by comparing the signal from the spiked biological sample (after subtracting endogenous levels from the unspiked sample) to the signal obtained from the spiked neat solution.

Second, since we added the same amount of IS in all samples, we also compared the IS peak areas in blood and brain samples to those in neat solution. Differences in IS peak area of less than 20% were considered acceptable. A decrease in IS peak area of more than 20% indicated ion suppression, while an increase of more than 20% indicated ion enhancement.

### Behavior and tissue collection for method application

We quantified 11 estrogens in wild male song sparrows. In the breeding season and non-breeding season, subjects were randomly assigned to one of two treatment groups: simulated territorial intrusion (STI) or control (CON) (n=10 subjects per group per season). Subjects in the STI group were exposed to a conspecific song playback from a speaker and a live caged decoy placed within their territory for 10 min. In contrast, subjects in the CON group were exposed to an empty cage and a silent speaker [26]. During the CON and STI conditions, aggressive behaviors were recorded [50,51], including song latency, flight latency, number of songs, number of flights, and time spent within 5 m of the decoy. Then, a mist net (set up in advance) was quickly unfurled and subjects were captured. The playback used for capture had similar duration across all four groups (breeding CON: 2.2 ± 0.6 min, breeding STI: 1.4 ± 0.6 min, non-breeding CON: 2.0 ± 0.4 min, non-breeding STI: 1.5 ± 0.4 min, F_3,36_ = 1.27, p = 0.30).

Immediately after capture, the subjects were euthanized within 3 min, and handling duration was similar across groups (breeding CON: 2.5 ± 0.1 min, breeding STI: 2.5 ± 0.2 min, non-breeding CON: 2.1 ± 0.1 min, non-breeding STI: 2.6 ± 0.1 min, F_3,36_ = 2.61, p = 0.066).

### Brain dissection

The Palkovits punch technique [52] was used to microdissect 9 brain regions that regulate social behavior (for details, see [26,45,49]. These regions include the caudal portion of the preoptic area (POA), anterior hypothalamus (AH), lateral septum (LS), bed nucleus of the stria terminalis (BNST), ventromedial hypothalamus (VMH), ventral tegmental area (VTA), central grey (CG), caudomedial nidopallium (NCM), and nucleus taeniae of the amygdala (TnA) (homolog of the mammalian medial amygdala). We also included the cerebellum (Cb), which has low aromatase expression and is not in the social behavior network. Brains were sectioned coronally at 300 μm using a cryostat, and bilateral punches were collected. The same punch size (1 mm diameter) was used for all regions. Depending on the size of the region, either 4 or 6 punches were collected, yielding 0.98 or 1.47mg of tissue, respectively. Punches were expelled into cold 2-mL polypropylene tubes (Sarstedt AG & Co, Numbrecht Germany), each containing five zirconium ceramic oxide beads (1.4-mm diameter) and stored at ‒70°C.

### Statistical analysis

A value was considered below the LLOQ if it fell below the lowest standard on the calibration curve. When 20% or more of the samples in a group (blood or brain region) were above the LLOQ, then the values below the LLOQ were imputed as before [26,45,49,53,54]. If less than 20% of the samples in a group were above the LLOQ, then imputations were not performed.

Statistical analyses were conducted using GraphPad Prism version 10.3.1 (GraphPad Software). When necessary, data were log-transformed prior to analysis. Seasonal differences in behavior were analyzed by unpaired t-tests. The correlations between steroid levels were examined using Spearman’s rho correlations. Regional differences in estrogen levels were analyzed using repeated measures one-way ANOVA followed by Tukey’s multiple comparison tests, with corrected p values reported. The significance criterion was set at p ≤ 0.05. Graphs show the mean ± SEM and are presented with non-transformed data.

## Results

### Assay development and validation

An assay for quantifying 11 estrogens was successfully developed using DMIS derivatization and LC-MS/MS. First, we achieved high specificity by identifying at least one MRM transition that is specific to each estrogen, and through the optimization of the liquid chromatography, which allowed the separation of isomers that shared MRM transitions (Table 1). Second, assay sensitivity was greatly enhanced for all estrogens after DMIS derivatization, as evidenced by the improved LLOQs (Table 2) and the linearity across the range (Fig. 1). Third, assay accuracy, as indicated by low and high QCs, generally fell within an acceptable range (100 ± 20%), with the exception of 4OH-E_2_ which was below 80% (Table 2). Fourth, assay precision, as indicated by intra- and inter-assay coefficients of variation, was considered acceptable when lower than 20%. The analytes E1, 17β-E2, E3, 2Me-E1, and 16α-OHE1 showed acceptable precision at both low and high QC concentrations (Table 2), for the rest of the analytes variability for the was higher than the desired for either low or high concentration QCs. Lastly, no peaks were detected in all blanks and double blanks for all analytes.

**Fig 1.**
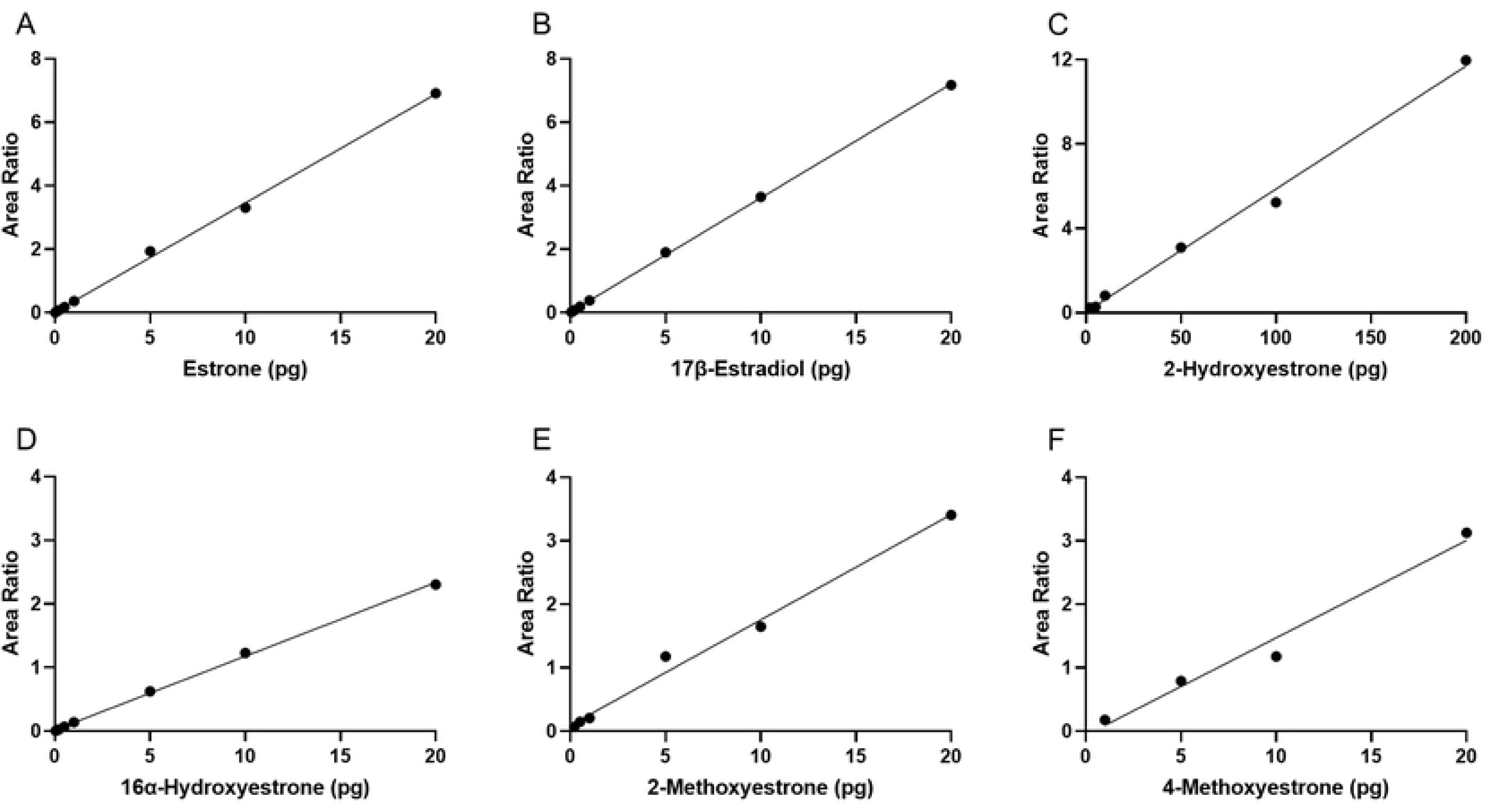
Calibration curves for 6 derivatized estrogen analytes measured by liquid chromatography-tandem mass spectrometry (LC-MS/MS). (A) estrone, (B) 17β-estradiol, (C) 2-hydroxyestrone, (D) 16α-hydroxyestrone, (E) 2-methoxyestrone, and (F) 4-methoxyestrone. The calibration curve for 2-hydroxyestrone is one order of magnitude higher than the other analytes, because of lower sensitivity for this analyte. Area ratio is calculated by dividing an analyte peak area by the appropriate internal standard (IS) peak area in the same sample.

**Table 2.** Assay lower limits of quantification (LLOQ), accuracy, and precision.

|  |  | Low concentration Quality Control |  |  | High concentration Quality Control |  |  |
| --- | --- | --- | --- | --- | --- | --- | --- |
| Analyte | LLOQ<br>(pg) | intra-assay<br>CV% | inter-assay<br>CV% | accuracy<br>% | intra-assay<br>CV% | inter-assay<br>CV% | accuracy<br>% |
| E<sub>1</sub> | 0.02 | 9.1 | 16.1 | 84.9 | 7.3 | 12.4 | 86.3 |
| 17β-E<sub>2</sub> | 0.05 | 3.9 | 6.0 | 92.8 | 2.8 | 3.7 | 86.1 |
| 17α-E<sub>2</sub> | 0.1 | 13.6 | 22.2 | 90.1 | - | - | - |
| E<sub>3</sub> | 0.02 | 2.0 | 3.0 | 87.5 | 1.9 | 2.5 | 86.6 |
| 2OH-E<sub>1</sub> | 2 | 15.9 | 28.7 | 89.7 | 10.0 | 24.7 | 106.7 |
| 2Me-E<sub>1</sub> | 0.2 | 13.3 | 20.6 | 97.5 | 13.4 | 15.5 | 94.5 |
| 4Me-E<sub>1</sub> | 1 | - | - | 95.2 | 29.8 | 37.0 | 92.5 |
| **16α-OHE<sub>1</sub>** | 0.05 | 3.7 | 7.8 | 85.7 | 3.3 | 5.5 | 84.2 |
| **4OH-E<sub>2</sub>** | 2 | 14.8 | 27.2 | 76.1 | 23.2 | 28.5 | 105.8 |
| **2Me-E<sub>2</sub>** | 0.2 | 17.4 | 24.8 | 96.7 | 11.8 | 18.4 | 91.3 |
| **4Me-E<sub>2</sub>** | 0.5 | 19.0 | 25.2 | 101.3 | 16.8 | 19.1 | 96.2 |

Note. The lower limit of quantification (LLOQ) was defined as the lowest standard on the calibration
curve with a signal-to-noise ratio greater than 10. The low quality control (QC) was 0.5 pg for each
analyte, except for 2OH-E<sub>1</sub> and 4OH-E<sub>2</sub> (5 pg). The high QC was 2 pg, except for 2OH-E<sub>1</sub> and 4OH-E<sub>2</sub>
(50 pg). For 4Me-E<sub>1</sub> the low QC was below its LLOQ. For 17α-E<sub>2</sub> only the low QC was used.

Four ^13^C-labeled estrogens were used as ISs. In general, ^13^C-labeled ISs are preferred over deuterated ISs because ^13^C-labeled IS are more stable and their retention times more closely match those of the analytes (Fig 2). One ^13^C-labeled estrogen was used for each group of estrogens. ^13^C_3_-E_3_ was used for E_3_ and 16α-OHE_1_; ^13^C_6_-E_2_ was used for E_1_, 17β-E_2_, and 17α-E_2_; ^13^C_6_-2Me-E_2_ was used for 2Me-E_1_, 4Me-E_1_, 2Me-E_2_, and 4Me-E_2_; and ^13^C_6_-2OH-E_2_ was used for 2OHE_1_ and 4OH-E_2_.

**Fig 2.**
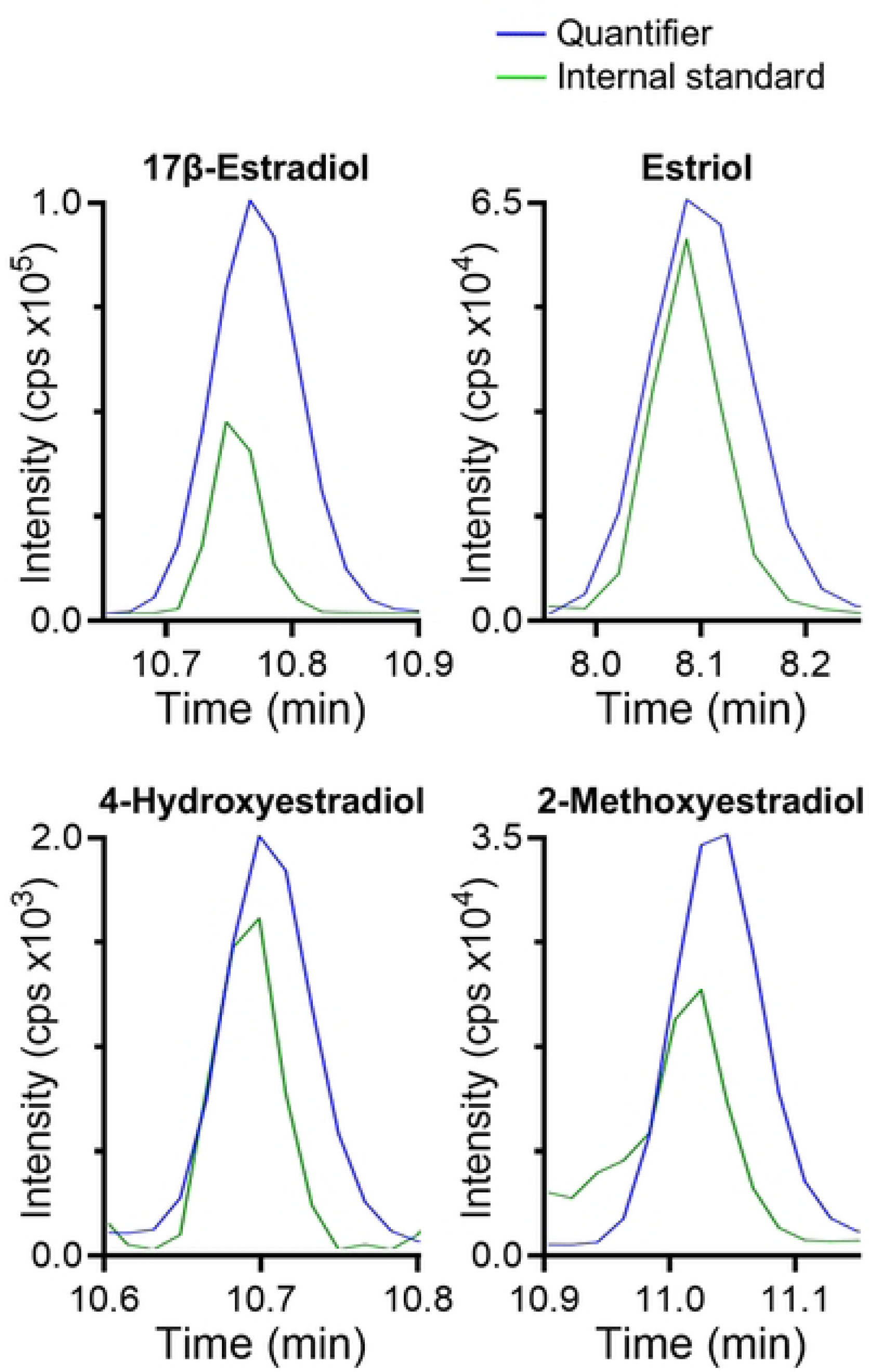
Representative chromatograms for 4 derivatized estrogen analytes measured by liquid chromatography-tandem mass spectrometry (LC-MS/MS). The samples were in neat solution for (A) 17β-estradiol (17β-E_2_), (B) estriol (E_3_), (C) 4-hydroxyestradiol (4OH-E_2_), and (D) 2-methoxyestradiol (2Me-E_2_). The quantifier transitions of the reference standards are shown in blue (10 pg of each) and the ^13^C-labeled internal standard transitions are shown in green (2 pg of each). Intensity is measured in counts per second (cps).

Matrix effects were assessed in blood (20 µL) and brain samples (1.5 mg). A total of n=5 subjects were used for method development. To assess recovery, we compared estrogen levels in samples spiked with known amounts of standard (after subtracting the endogenous concentrations of unspiked tissue samples) to those of standards spiked in a neat solution.

Recovery percentages within the range of 100 ± 20% are considered acceptable. Recovery values for the 11 analytes are presented in Table 3. For E_1_, 17β-E_2_, E_3_, and 16α-OHE_1_ recovery values were within the acceptable range. The catecholestrogens, 2OHE_1_ and 4OH-E_2_, showed acceptable recoveries in brain but lower recoveries in blood. Methoxyestrogens showed variable performance: 4Me-E_2_ had good recovery; 4Me-E_1_ showed low recovery in brain; 2Me-E_1_ and 2Me-E_2_ showed affected recovery in both brain and blood.

**Table 3.** Assay recovery %.

|  | **Blood (20μL)** | **Brain (1.5 mg)** |
| --- | --- | --- |
| **E<sub>1</sub>** | 99 | 106 |
| **17β-E<sub>2</sub>** | 91 | 105 |
| **E<sub>3</sub>** | 105 | 105 |
| **2OH-E<sub>1</sub>** | 48 | 86 |
| **2Me-E<sub>1</sub>** | 211 | 126 |
| **4Me-E<sub>1</sub>** | 88 | 76 |
| **16α-OHE<sub>1</sub>** | 118 | 104 |
| **4OH-E<sub>2</sub>** | 25 | 113 |
| **2Me-E<sub>2</sub>** | 74 | 130 |
| **4Me-E<sub>2</sub>** | 84 | 82 |

Matrix effects were also assessed by comparing IS peak areas in blood and brain samples to those in neat solution (Table 4). Tissue IS peak areas were considered acceptable in a range of 100 ± 20%. A decrease of more than 20% indicates ion suppression, while an increase of more than 20% indicates ion enhancement. The ^13^C_6_-E_2_ IS showed comparable peak areas in both blood and brain relative to the neat solution, suggesting minimal matrix effects. The ^13^C_3_-E_3_ IS exhibited potential ion suppression in blood but remained within the acceptable range in brain tissue. In contrast, ^13^C_6_-2OH-E_2_ and ^13^C_6_-2Me-E_2_ showed higher IS peak areas in both biological matrices, indicating possible ion enhancement. Nonetheless, catecholestrogens and methoxyestrogens analytes demonstrated better recovery performance than their respective IS (Table 3), indicating that the ISs partially corrected for matrix effects.

**Table 4.** Internal standard peak area in each matrix relative to neat solution (%)

|  | **Blood (20μL)** | **Brain (1.5 mg)** |
| --- | --- | --- |
| $^{13}\text{C}_6\text{-E}_2$ | 87 | 94 |
| $^{13}\text{C}_3\text{-E}_3$ | 62 | 97 |
| $^{13}\text{C}_6\text{-2OH-E}_2$ | 812 | 422 |
| $^{13}\text{C}_6\text{-2Me-E}_2$ | 260 | 188 |

### Method application in songbird samples

As expected, male song sparrows responded aggressively to the STI during both the breeding and non-breeding seasons, and no significant seasonal differences were found in any of the behaviors assessed (Table 5). These data indicate that the sparrows respond with similar levels of aggression to an intruder during these two seasons, as in previous studies [55,56].

**Table 5.**
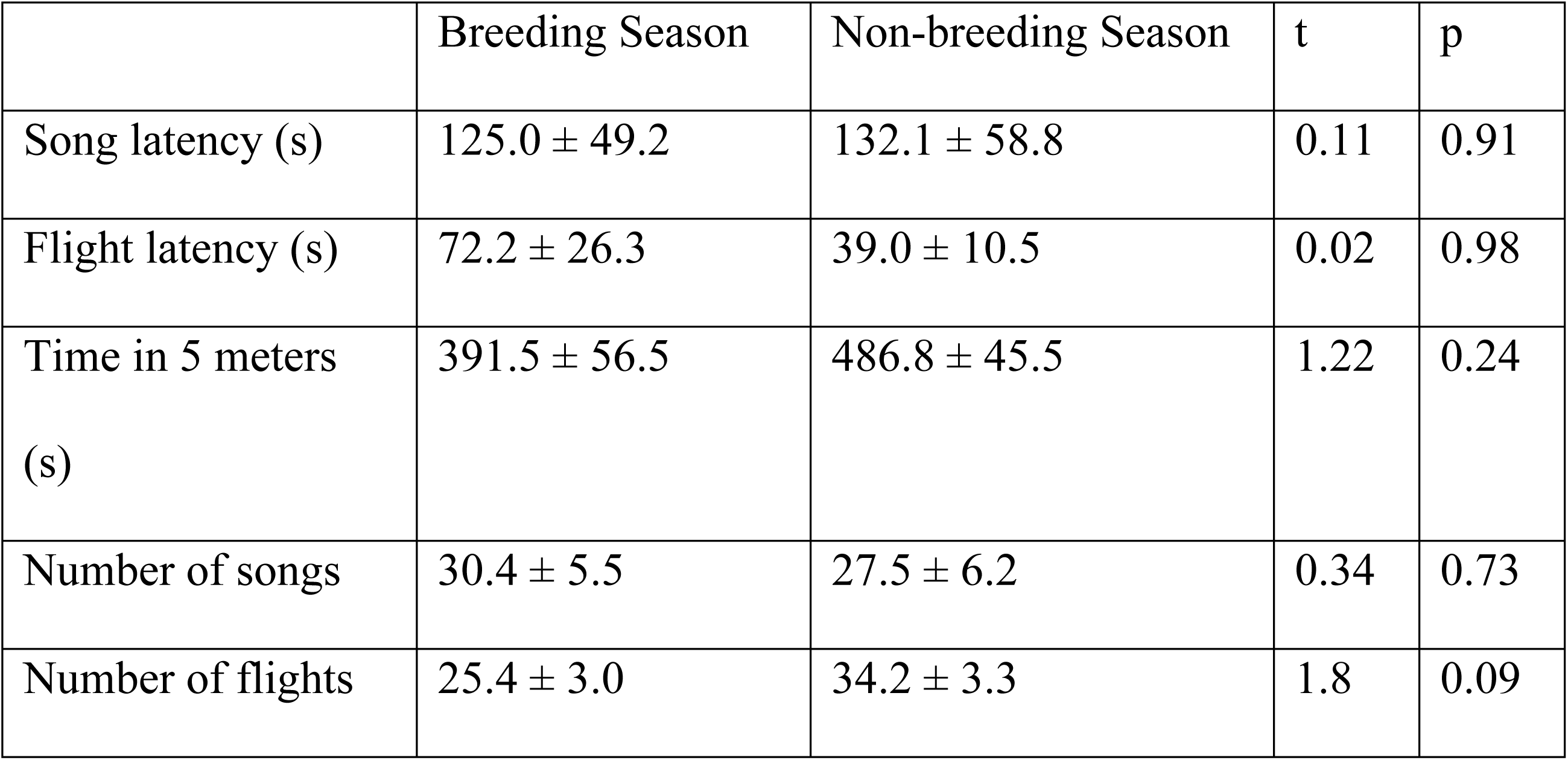
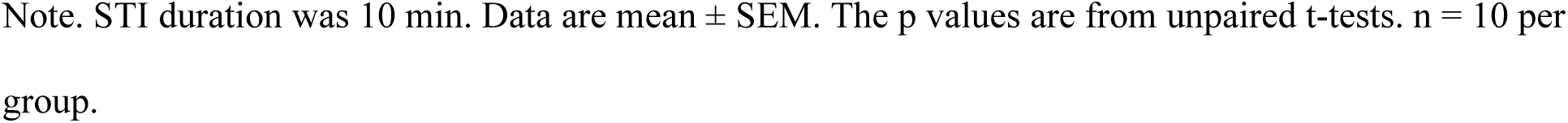
Behavioral responses to a Simulated Territorial Intrusion (STI) in wild song sparrows.

|  | Breeding Season | Non-breeding Season | t | p |
| --- | --- | --- | --- | --- |
| Song latency (s) | 125.0 ± 49.2 | 132.1 ± 58.8 | 0.11 | 0.91 |
| Flight latency (s) | 72.2 ± 26.3 | 39.0 ± 10.5 | 0.02 | 0.98 |
| Time in 5 meters (s) | 391.5 ± 56.5 | 486.8 ± 45.5 | 1.22 | 0.24 |
| Number of songs | 30.4 ± 5.5 | 27.5 ± 6.2 | 0.34 | 0.73 |
| Number of flights | 25.4 ± 3.0 | 34.2 ± 3.3 | 1.8 | 0.09 |

Note. STI duration was 10 min. Data are mean ± SEM. The p values are from unpaired t-tests. n = 10 per group.

We applied the validated method to examine a panel of 11 estrogens in the blood and 10 microdissected brain regions of CON and STI subjects in the two seasons (n=10 per treatment per season). This generated 121 data points per subject, and thus 4,840 data points across 40 subjects total.

In the breeding season, only E_1_ and 17β-E_2_ were quantifiable in blood and brain samples, in both the CON and STI groups (Fig. 3). For E_1_, STI had no significant effect on blood levels (t = 0.76, p = 0.46). In the brain, there was a significant main effect of region on E_1_ levels (F(9,162) = 39.05; p < 0.0001) but no significant main effect of STI (F(1,18) = 0.25; p = 0.63) and no significant region × STI interaction (F(9,162) = 0.76; p = 0.65). Similarly, for 17β-E_2_, levels in blood were not significantly different between CON and STI groups, although there was a trend for STI to increase 17β-E_2_ levels (t = 2.01, p = 0.06). In the brain, 17β-E_2_ levels showed a significant main effect of region (F(9,162) = 166.5; p < 0.0001) but no significant main effect of STI (F(1,18) = 0.0014; p = 0.97) and no significant region × STI interaction (F(9,162) = 0.76; p = 0.65).

**Fig 3.**
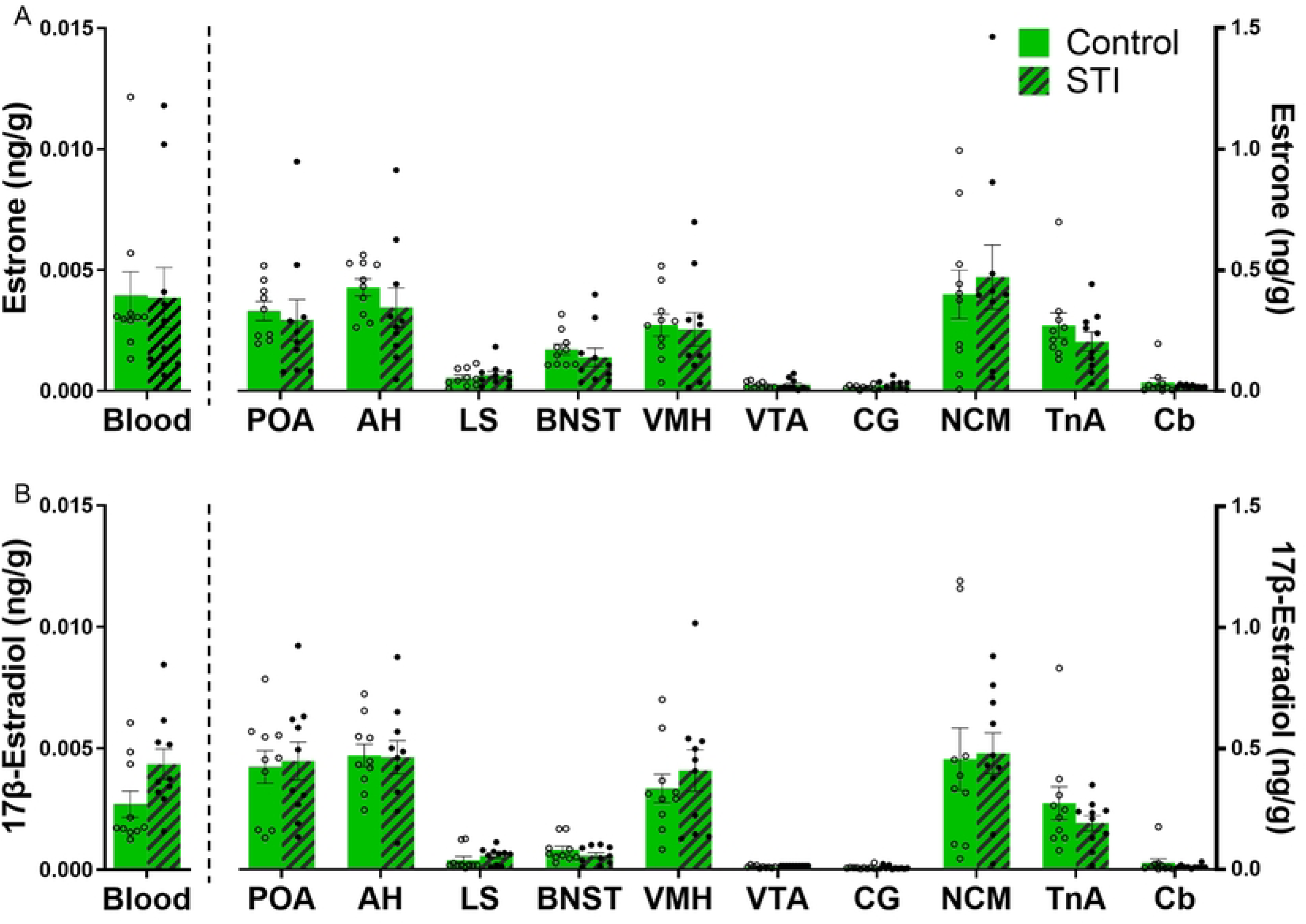
Lack of effect of a 10-min STI on circulating and brain estrogen levels of wild adult male song sparrows during the breeding season. The bar graphs represent concentrations of (A) E_1_ and (B) 17β-E_2_. Values are expressed as the mean ± SEM. n = 10 per group.

In the non-breeding season, neither E_1_ nor 17β-E_2_ were quantifiable in any samples from both groups.

The other nine estrogens in the panel were below the LLOQs in blood and brain in both groups and both seasons.

Next, in samples from the breeding season, we assessed the relationships between 17β-E_2_and its precursor E_1_ in each brain area. As expected, there were significant positive correlations between E_1_ and 17β-E_2_ in each group separately (data not shown) and both groups combined (Fig. 4) in all brain areas, except the VTA where no correlation was observed, probably due to its low estrogen levels.

**Fig 4.**
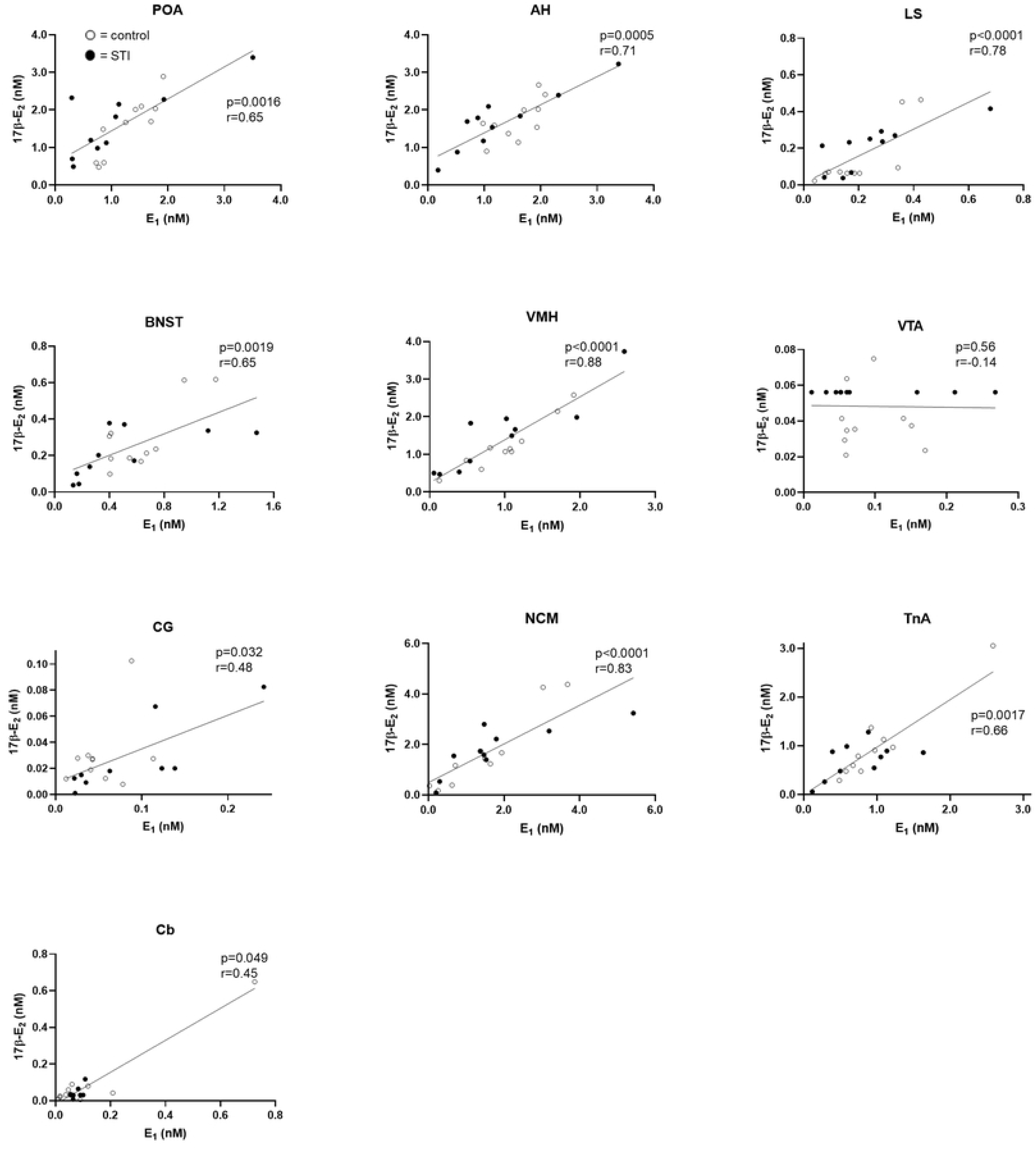
Correlations between estrogens in each brain area in wild adult male song sparrows during the breeding season. Levels of E_1_ and 17β-E_2_ were significantly positively correlated in all brain regions except the VTA. Data were analyzed using Spearman’s rho. Subjects from CON and STI groups are represented with empty and filled circles, respectively. n = 10 per group.

## Discussion

We attempted to develop a quantification method for 11 estrogens, adding 4 new analytes and 4 new ^13^C-labeled internal standards to our previous method [45]. Overall, assay performance was satisfactory for E_1_, 17α-E_2_, 17β-E_2_, E_3_, 4Me-E_1_, 16α-OHE_1_, and 4Me-E_2_ in blood and plasma. For the methoxyestrogens, 2Me-E_1_ and 2Me-E_2_, performance was hindered in both matrices. Assay performance for catecholestrogens was acceptable in brain tissue, but matrix effects were observed for these analytes in blood. The improvements presented here are, in part, the result of the four ^13^C-labeled IS that we used. We then applied the method to wild song sparrows and examined the effects of tissue, season, and an aggressive encounter. We detected only E_1_ and 17β-E_2_. The tissue differences were robust, and E_1_ and 17β-E_2_ levels were higher in brain regions that express aromatase. The seasonal effect was also robust, and estrogens were higher during the breeding season than the non-breeding season. Male song sparrows showed intense aggression during both seasons, yet the aggressive encounter did not acutely affect the concentration of E_1_ and 17β-E_2_ in brain or blood.

### Method development and validation

Estrogen measurement is particularly challenging due in part to low abundance of estrogens in biological samples, but mass spectrometry assay sensitivity can be improved through derivatization [38,39,41,57], as seen here. We separated analytes by ultra-high-performance liquid chromatography (UHPLC), and each analyte exhibited a unique retention time, supporting both analyte identification and method specificity. Here, we used electrospray ionization (ESI). While atmospheric pressure photoionization is also used with DMIS [48], ESI is more widely employed. The DMIS derivatization reagent produced analyte-specific fragmentation patterns, a further contributing to assay specificity. This contrasts with dansyl chloride, the most commonly used reagent for 17β-E_2_ derivatization, which generates a product ion derived from the dansyl moiety itself, and thus is not analyte-specific [58,59]. Our approach therefore represents a methodological improvement by providing enhanced specificity for estrogen detection.

Importantly, most studies of estrogens exclusively measure 17β-E_2_, and the levels of other estrogens are under-investigated. Both E_1_ and 17β-E_2_ undergo hydroxylation at the C2 and C4 positions, generating bioactive catecholestrogens [60,61]. Although relatively labile [62], these compounds have diverse reproductive [63,64], neuromodulatory [65,66], and anti- or pro-carcinogenic effects [67]. These catecholestrogens are converted into methoxyestrogens, which are subsequently conjugated to glucuronide or sulfate groups for excretion [61,68,69]. In this study, we expanded our previous method [45] by incorporating 4 additional estrogens (16α-OHE_1_, 2OH-E_1_, 2Me-E_1_, and 4Me-E_1_) to our original panel of E_1_, 17β-E_2_, 17α-E_2_, E_3_, 4OH-E_2_, 2Me-E_2_, and 4Me-E_2_. Accuracy was acceptable for all analytes, even at the lower concentrations. While greater variability was observed for the catechol- and methoxy-estrogens, assay precision was acceptable overall. The assay demonstrated excellent sensitivity, as low as 0.02 pg per sample, making this method well-suited for quantifying neuroestrogens in microdissected brain regions, ranging from 0.01 to 2.5 pg per sample [45,49,70], and thus below the LLOQs of most assays.

In the present study we further improved our estrogen quantification method by incorporating four ^13^C-labeled ISs. While most studies use deuterated ISs, which are more widely available and affordable, these compounds are more susceptible to hydrogen-deuterium exchange during sample processing, potentially compromising assay accuracy [31]. Additionally, deuterated ISs often exhibit slightly different retention times than their corresponding analytes.

In contrast, ^13^C-labeled IS more closely match the retention times of their analytes, resulting in improved co-elution and enhanced assay performance [39]. Moreover, in our previous study, we used a single deuterated IS (d4-17β-E_2_) for all seven estrogen analytes [45]. In contrast, in the present study, we used four stable-isotope labelled IS, with one labelled estrogen per chemical group. This refinement improved matrix effects particularly for catechol- and methoxy-estrogens. For example, recovery of 4OH-E_2_ in our previous study was 613% in brain, but 113% here (similar results are observed for both 2Me-E_2_ and 4Me-E_2_). Thus, although some matrix effects were observed here, we notably improved analyte recovery compared to our previous method, in which catechol- and methoxy-estrogens were strongly affected by matrix effects. Furthermore, the assay showed better performance in brain tissue, highlighting its suitability for neurosteroid analysis.

Further evidence supports our assay performance. Estrogen levels detected in the present study were similar to those found in previous studies in this species, either with [45] or without [26,49] derivatization. Moreover, we observed a positive correlation between E_1_ and 17β-E_2_ across nearly all brain regions examined, as expected. DMIS derivatization has been applied to mouse and human plasma [44,48,71], and human saliva [72], demonstrating its potential applications in other species and sample types.

### Method application in wild songbirds

We applied the assay to wild song sparrows to examine how tissue type, season, and an aggressive encounter affect estrogen levels. In the non-breeding season, estrogens were non-detectable in both CON and STI subjects. In contrast, in the breeding season, both E_1_ and 17β-E_2_ were above the LLOQ in the brain in both groups, at higher levels than in circulation, indicating robust tissue-specific differences. Additionally, E_1_ and 17β-E_2_ showed pronounced regional variation across the brain, matching known regional differences in aromatase. All of these patterns are consistent with our previous findings in song sparrows under baseline conditions [26,45,49]. However, we found no effect of the aggressive challenge on E_1_ and 17β-E_2_ levels in either brain or blood, regardless of season.

Neuroestrogens promote aggression in non-breeding song sparrows, yet we found no evidence of their synthesis during this season. In the song sparrow brain, aromatase is highly expressed year-round [19,27]. Interestingly, in the non-breeding season only, acute aromatase inhibition reduces aggression [28,29] and exogenous 17β-E_2_ rapidly increases aggressive behavior [30]. Our previous study found an increase in brain androstenedione and testosterone within 10 min of aggressive interactions [26], which could provide the substrate for aromatase to produce neuroestrogens during contests. We therefore predicted that a territorial challenge would elevate brain estrogens in both seasons. However, this was not observed. In the breeding season, E_1_ and 17β-E_2_ were present but unchanged after STI, and in the non-breeding season, brain estrogens were non-detectable in both groups.

One possible explanation is that our sampling time might have missed the peak of estrogen production. Subjects were sampled after 10 min of aggressive interaction, which may have been either too early or too late. Since aromatase is present in the song sparrow brain, aromatase activation may rely on post-translational modifications such as phosphorylation [73,74] to rapidly produce the neuroestrogens. In quail, increased aromatase activity can occur as early as 2 min after aggressive interactions (whose activity also correlates with aggression levels) [10], and is similarly modulated within 1–5 minutes during sexual encounters, returning to baseline by 15 min [22]. Alternatively, it could be that estrogens take longer times to increase, and we collected tissues too early. In white-crowned sparrows and zebra finches, local 17β-E_2_ levels and aromatase activity are modulated 30 min after a social interaction [23,25,75]. In song sparrows, studies employing acute pharmacological modulations have used longer timeframes, ranging from 20 min to 24 hr [29,30].

A second possibility is that we did not measure the right analytes. Estrogens are a large and diverse family of compounds, and our method might have omitted critical compounds. For instance, estrogens can be conjugated to glucuronide or sulfate moieties [61]. Neuroestrogens might be synthesized and then promptly converted to these conjugated forms, which we did not measure.

Third, the estrogenic modulation of non-breeding aggression could be due to changes in estrogen sensitivity. Estrogen receptors are expressed in the song sparrow brain during both breeding and non-breeding seasons with no seasonal differences [27]. Interestingly, receptor availability can be modulated by trafficking to and from the cell membrane [76], and receptor function can be rapidly modulated by phosphorylation [77], potentially altering sensitivity to estrogens without requiring changes in ligand concentration. However, this is unlikely due to the changes in behavior observed in aromatase enzyme manipulation experiments.

Lastly, perhaps our stimuli were insufficient to trigger a neuroendocrine response. Subjects were exposed to a live decoy in a transparent cage accompanied by a conspecific song playback. Although this stimulus reliably elicited a robust behavioral aggressive response, it may not have provided the full suite of cues necessary to activate neural estrogen synthesis. For example, studies in quail showed that males exhibit different neuroestrogen patterns depending on whether they have visual access only or full access to females, with copulation producing the strongest effects [78]. Moreover, in our paradigm, the decoy did not directly interact with the subject. In song sparrows, endocrine responses, such as testosterone elevation, depend on the sensory modality of the social cue, with combined visual and auditory stimuli being more effective than either alone [79]. However, previous studies have shown that aromatase inhibition still reduces aggressive responses to this same STI setup [28,29], suggesting that neural estrogens are indeed relevant in this context, even if we were unable to detect them here.

## Conclusions

We developed and validated a highly sensitive LC-MS/MS method for the simultaneous quantification of eleven estrogens in microdissected songbird blood and brain. This approach significantly improves upon previous methods, by reducing matrix effects and incorporating additional analytes to the panel. Application of the method to free-living male song sparrows exposed to an acute social challenge did not show changes in neuroestrogen levels, despite a strong behavioral response. Our refined method offers a powerful tool for quantifying estrogens in brain, with potential applicability across diverse vertebrate systems.

## Acknowledgments

We thank Asmita Poudel for her assistance with LC-MS/MS and Sofia Gray for help with fieldwork. Brittany Jensen, Sofia Laforest, and Guillermo Valiño for comments on the manuscript.

## References

1. Shanmugan S, Epperson CN. Estrogen and the prefrontal cortex: Towards a new understanding of estrogen’s effects on executive functions in the menopause transition. Hum Brain Mapp. 2014;35:847–65. doi: 10.1002/hbm.22218

2. Gervais NJ, Remage-Healey L, Starrett JR, Pollak DJ, Mong JA, Lacreuse A. Adverse effects of aromatase inhibition on the brain and behavior in a nonhuman primate. J Neurosci. 2019;39(5):918–28. doi: 10.1523/JNEUROSCI.0353-18.2018

3. Tinkler GP, Voytko M Lou. Estrogen modulates cognitive and cholinergic processes in surgically menopausal monkeys. Prog Neuro-Psychopharmacology Biol Psychiatry. 2005;29:423–31. doi: 10.1016/j.pnpbp.2004.12.016

4. Rosvall KA, Bentz AB, George EM. How research on female vertebrates contributes to an expanded challenge hypothesis. Horm Behav. 2020;123(April 2019):104565. doi: 10.1016/j.yhbeh.2019.104565

5. Quintana L, Jalabert C, Fokidis HB, Soma KK, Zubizarreta L. Neuroendocrine Mechanisms Underlying Non-breeding Aggression: Common Strategies Between Birds and Fish. Front Neural Circuits. 2021;15(July). doi: 10.3389/fncir.2021.716605

6. Jalabert C, Munley KM, Demas GE, Soma KK. Aggressive Behavior. In: Encyclopedia of Reproduction. second. Academic Press; 2018. p. 242–7. doi: 10.1016/B978-0-12-801238-3.64591-9

7. Plumier JP, Jalabert C, Munley KM, Demas GE, Soma KK. Aggressive Behavior. In: Reference Module in Biomedical Sciences. Elsevier; 2024. p. 242–7. doi: 10.1016/B978-0-443-21477-6.00303-5

8. Trainor BC, Sima Finy M, Nelson RJ. Rapid effects of estradiol on male aggression depend on photoperiod in reproductively non-responsive mice. Horm Behav. 2008;53(1):192–9. doi: 10.1016/j.yhbeh.2007.09.016

9. Trainor BC, Lin S, Finy* MS, Rowland MR, Nelson RJ. Photoperiod reverses the effects of estrogens on male aggression via genomic and nongenomic pathways. Proc Natl Acad Sci. 2007;104(23):9840–5. doi: 10.1073/pnas.0701819104

10. Schlinger BA, Callard G V. Aromatization mediates aggressive behavior in quail. Gen Comp Endocrinol. 1990;79(1):39–53. doi: 10.1016/0016-6480(90)90086-2

11. Huffman LS, O’Connell LA, Hofmann HA, O’Connell LA, Hofmann HA. Aromatase regulates aggression in the African cichlid fish Astatotilapia burtoni. Physiol Behav. 2013;112–113(0):77–83. doi: 10.1016/j.physbeh.2013.02.004

12. Merritt JR, Davis MT, Jalabert C, Libecap TJ, Williams DR, Soma KK, et al. Rapid effects of estradiol on aggression depend on genotype in a species with an estrogen receptor polymorphism. Horm Behav. 2018;98:210–8. doi: 10.1016/j.yhbeh.2017.11.014

13. Scaia MF, Morandini L, Noguera CA, Trudeau VL, Somoza GM, Pandolfi M. Can estrogens be considered as key elements of the challenge hypothesis? The case of intrasexual aggression in a cichlid fish. Physiol Behav. 2018;194(February):481–90. doi: 10.1016/j.physbeh.2018.06.028

14. Maney DL, Goode CT, Lange HS, Sanford SE, Solomon BL. Estradiol modulates neural responses to song in a seasonal songbird. J Comp Neurol. 2008;511:173–86. doi: 10.1002/cne.21830

15. Remage-Healey L, Jeon SD, Joshi NR. Recent evidence for rapid synthesis and action of estrogens during auditory processing in a song. J Neuroendocrinol. 2013;25(11):1024–31. doi: 10.1111/jne.12055

16. Yoder KM, Vicario DS. To Modulate and Be Modulated: Estrogenic Influences on Auditory Processing of Communication Signals within a Socio-Neuro-Endocrine Framework. Behav Neurosci. 2012;126(1):17–28. doi: 10.1037/a0026673

17. George EM, Rosvall KA. How a territorial challenge changes sex steroid-related gene networks in the female brain: A field experiment with the tree swallow. Horm Behav. 2025;169:105698. doi: 10.1016/j.yhbeh.2025.105698

18. Shen P, Schlinger BA, Campagnoni AT, Arnold AP. An atlas of aromatase mRNA expression in the zebra finch brain. J Comp Neurol. 1995;360(1):172–84. doi: 10.1002/cne.903600113

19. Soma KK, Schlinger BA, Wingfield JC, Saldanha CJ. Brain aromatase, 5α-reductase, and 5β-reductase change seasonally in wild male song sparrows: Relationship to aggressive and sexual behavior. J Neurobiol. 2003;56(3):209–21. doi: 10.1002/neu.10225

20. Forlano PM, Schlinger BA, Bass AH. Brain aromatase: New lessons from non-mammalian model systems. Front Neuroendocrinol. 2006;27(3):247–74. doi: 10.1016/j.yfrne.2006.05.002

21. Roselli CE. The Distribution and Regulation of Aromatase in the Mammalian Brain: from Mice to Monkeys. In: Balthazart J, Ball G, editors. Brain Aromatase, Estrogens, and Behavior. Oxford Academic; 2012. p. 43–63. doi: 10.1093/acprof:oso/9780199841196.001.0001

22. Cornil CA, Dalla C, Papadopoulou-Daifoti Z, Baillien M, Dejace C, Ball GF, et al. Rapid decreases in preoptic aromatase activity and brain monoamine concentrations after engaging in male sexual behavior. Endocrinology. 2005;146(9):3809–20. doi: 10.1210/en.2005-0441

23. Charlier TD, Newman AEM, Heimovics SA, Po KWL, Saldanha CJ, Soma KK. Rapid Effects of Aggressive Interactions on Aromatase Activity and Oestradiol in Discrete Brain Regions of Wild Male White-Crowned Sparrows. J Neuroendocrinol. 2011;23(8):742–53. doi: 10.1111/j.1365-2826.2011.02170.x

24. Rosvall KA, Bergeon Burns CM, Barske J, Goodson JL, Schlinger BA, Sengelaub DR, et al. Neural sensitivity to sex steroids predicts individual differences in aggression: Implications for behavioural evolution. Proc R Soc B Biol Sci. 2012;279(1742):3547–55. doi: 10.1098/rspb.2012.0442

25. Remage-Healey L, Maidment NT, Schlinger BA. Forebrain steroid levels fluctuate rapidly during social interactions. Nat Neurosci. 2008;11(11):1327–34. doi: 10.1038/nn.2200.Forebrain

26. Jalabert C, Gray SL, Soma KK. An aggressive interaction rapidly increases brain androgens in a male songbird during the non-breeding season. J Neurosci. 2024;e1095232024. doi: 10.1523/JNEUROSCI.1095-23.2024

27. Wacker DW, Wingfield JC, Davis JE, Meddle SL. Seasonal Changes in Aromatase and Androgen Receptor, but not Estrogen Receptor mRNA Expression in the Brain of the Free-Living Male Song Sparrow, Melospiza melodia morphna. J Comp Neurol. 2010;518(18):3819–35. doi: 10.1002/cne.22426

28. Soma KK, Tramontin AD, Wingfield JC. Oestrogen regulates male aggression in the non-breeding season. Proc R Soc B Biol Sci. 2000;267(1448):1089–96. doi: 10.1098/rspb.2000.1113

29. Soma KK, Sullivan KA, Tramontin AD, Saldanha CJ, Schlinger BA, Wingfield JC. Acute and chronic effects of an aromatase inhibitor on territorial aggression in breeding and nonbreeding male song sparrows. J Comp Physiol - A Sensory, Neural, Behav Physiol. 2000;186(7–8):759–69. doi: 10.1007/s003590000129

30. Heimovics SA, Ferris JK, Soma KK. Non-invasive administration of 17β-estradiol rapidly increases aggressive behavior in non-breeding, but not breeding, male song sparrows. Horm Behav. 2015;69:31–8. doi: 10.1016/j.yhbeh.2014.11.012

31. Wudy SA, Schuler G, Sánchez-guijo A, Hartmann MF. The art of measuring steroids Principles and practice of current hormonal steroid analysis. J Steroid Biochem Mol Biol. 2018;179:88–103. doi: 10.1016/j.jsbmb.2017.09.003

32. Handelsman DJ, Jones G, Kouzios D, Desai R. Evaluation of testosterone, estradiol and progesterone immunoassay calibrators by liquid chromatography mass spectrometry. Clin Chem Lab Med. 2023;61(9):1612–8. doi: 10.1515/cclm-2022-1179

33. Stanczyk FZ, Xu X, Sluss PM, Brinton LA, McGlynn KA. Do metabolites account for higher serum steroid hormone levels measured by RIA compared to mass spectrometry? Clin Chim Acta. 2018;484:223–5. doi: 10.1016/j.cca.2018.05.054

34. Haisenleder DJ, Schoenfelder AH, Marcinko ES, Geddis LM, Marshall JC. Estimation of estradiol in mouse serum samples: Evaluation of commercial estradiol immunoassays. Endocrinology. 2011;152(11):4443–7. doi: 10.1210/en.2011-1501

35. Faupel-Badger JM, Fuhrman BJ, Xu X, Falk RT, Keefer LK, Veenstra TD, et al. Comparison of liquid chromatography-mass spectrometry, radioimmunoassay, and enzyme-linked immunosorbent assay methods for measurement of urinary estrogens. Cancer Epidemiol biomarkers Prev. 2010;19(1):292–300. doi: doi:10.1158/1055-9965.EPI-09-0643

36. Handelsman DJ, Newman JD, Jimenez M, McLachlan R, Sartorius G, Jones GRD. Performance of direct estradiol immunoassays with human male serum samples. Clin Chem. 2014;60(3):510–7. doi: 10.1373/clinchem.2013.213363

37. Wang C, Catlin DH, Demers LM, Starcevic B, Swerdloff RS. Measurement of Total Serum Testosterone in Adult Men: Comparison of Current Laboratory Methods Versus Liquid Chromatography-Tandem Mass Spectrometry. J Clin Endocrinol Metab. 2004;89(2):534–43. doi: 10.1210/jc.2003-031287

38. Blair IA. Analysis of estrogens in serum and plasma from postmenopausal women: Past present, and future. Steroids. 2010;75(4–5):297–306. doi: 10.1016/j.steroids.2010.01.012

39. Denver N, Khan S, Homer NZM, MacLean MR, Andrew R. Current strategies for quantification of estrogens in clinical research. J Steroid Biochem Mol Biol. 2019;192(April):105373. doi: 10.1016/j.jsbmb.2019.04.022

40. Wang Q, Mesaros C, Blair IA. Ultra-high sensitivity analysis of estrogens for special populations in serum and plasma by liquid chromatography–mass spectrometry: Assay considerations and suggested practices. J Steroid Biochem Mol Biol. 2016;162:70–9. doi: 10.1016/j.jsbmb.2016.01.002

41. Andrew R, Homer NZM. Mass spectrometry: Future opportunities for profiling and imaging steroids and steroid metabolites. Curr Opin Endocr Metab Res. 2020;15:71–8. doi: 10.1016/j.coemr.2020.11.005

42. Faqehi AMM, Cobice DF, Naredo G, Mak TCS, Upreti R, Gibb FW, et al. Derivatization of estrogens enhances specificity and sensitivity of analysis of human plasma and serum by liquid chromatography tandem mass spectrometry. Talanta. 2016;151:148–56. doi: 10.1016/j.talanta.2015.12.062

43. Denver N, Khan S, Stasinopoulos I, Church C, Homer NZ, MacLean MR, et al. Derivatization enhances analysis of estrogens and their bioactive metabolites in human plasma by liquid chromatography tandem mass spectrometry. Anal Chim Acta. 2019;1054:84–94. doi: 10.1016/j.aca.2018.12.023

44. Handelsman DJ, Gibson E, Davis S, Golebiowski B, Walters KA, Desai R. Ultrasensitive serum estradiol measurement by liquid chromatography-mass spectrometry in postmenopausal women and mice. J Endocr Soc. 2020;4(9):1–12. doi: 10.1210/jendso/bvaa086

45. Jalabert C, Shock MA, Ma C, Bootsma TJ, Liu MQ, Soma KK. Ultrasensitive Quantification of Multiple Estrogens in Songbird Blood and Microdissected Brain by LC-MS/MS. eNeuro. 2022;9(4):1–16. doi: 10.1523/eneuro.0037-22.2022

46. Wang Q, Rangiah K, Mesaros C, Snyder NW, Vachani A, Song H, et al. Ultrasensitive quantification of serum estrogens in postmenopausal women and older men by liquid chromatography-tandem mass spectrometry. Steroids. 2015;96:140–52. doi: 10.1016/j.steroids.2015.01.014

47. Anari MR, Bakhtiar R, Zhu B, Huskey S, Franklin RB, Evans DC. Derivatization of ethinylestradiol with dansyl chloride to enhance electrospray ionization: Application in trace analysis of ethinylestradiol in rhesus monkey plasma. Anal Chem. 2002;74(16):4136–44. doi: 10.1021/ac025712h

48. Keski-Rahkonen P, Desai R, Jimenez M, Harwood DT, Handelsman DJ. Measurement of Estradiol in Human Serum by LC-MS/MS Using a Novel Estrogen-Specific Derivatization Reagent. Anal Chem. 2015;87(14):7180–6. doi: 10.1021/acs.analchem.5b01042

49. Jalabert C, Ma C, Soma KK. Profiling of systemic and brain steroids in male songbirds: Seasonal changes in neurosteroids. J Neuroendocrinol. 2021;33(1):1–16. doi: 10.1111/jne.12922

50. Wingfield JC. Short-term changes in plasma levels of hormones during establishment and defense of a breeding territory in male song sparrows, Melospiza melodia. Horm Behav. 1985;19(2):174–87. doi: 10.1016/0018-506X(85)90017-0

51. Wingfield JC, Hahn TP. Testosterone and territorial behaviour in sedentary and migratory sparrows. Anim Behav. 1994;47(1):77–89. doi: 10.1006/anbe.1994.1009

52. Palkovits M. Isolated removal of hypothalamic or other brain nuclei of the rat. Brain Res. 1973;59:449–50. doi: 10.1016/0006-8993(73)90290-4

53. Tobiansky DJ, Kachkovski G V, Enos RT, Schmidt KL, Murphy EA, Soma KK. Sucrose consumption alters steroid and dopamine signalling in the female rat brain. J Endocrinol. 2020;245(2):231–46. doi: 10.1530/JOE-19-0386

54. Tobiansky DJ, Kachkovski G V, Enos RT, Schmidt KL, Murphy EA, Floresco SB, et al. Maternal sucrose consumption alters behaviour and steroids in adult rat offspring. J Endocrinol. 2021;251(3):161–80. doi: 10.1530/JOE-21-0166

55. Newman AEM, Soma KK. Aggressive interactions differentially modulate local and systemic levels of corticosterone and DHEA in a wild songbird. Horm Behav. 2011;60(4):389–96. doi: 10.1016/j.yhbeh.2011.07.007

56. Wingfield JC, Soma KK. Spring and Autumn Territoriality in Song Sparrows: Same Behavior, Different Mechanisms? 1 [Internet]. Vol. 42. 2002. doi: 10.1093/icb/42.1.11

57. Price HR, Jalabert C, Seib DR, Ma C, Lai D, Soma KK, et al. Measurement of Steroids in the Placenta, Maternal Serum, and Fetal Serum in Humans, Rats, and Mice: A Technical Note. Separations. 2023;10(4):1–12. doi: 10.3390/separations10040221

58. Xu L, Spink DC. Analysis of steroidal estrogens as pyridine-3-sulfonyl derivatives by liquid chromatography electrospray tandem mass spectrometry. Anal Biochem. 2008;375(1):105–14. doi: 10.1016/j.ab.2007.11.028

59. Li X, Franke AA. Improved profiling of estrogen metabolites by orbitrap LC/MS. Steroids. 2015;99:84–90. doi: 10.1016/j.steroids.2014.12.005

60. Sepkovic DW, Bradlow HL. Estrogen hydroxylation - The good and the bad. Ann N Y Acad Sci. 2009;1155:57–67. doi: 10.1111/j.1749-6632.2008.03675.x

61. Raftogianis R, Creveling C, Weinshilboum R, Weisz J. Estrogen metabolism by conjugation. J Natl Cancer Inst Monogr. 2000;27:113–24. doi: 10.1093/oxfordjournals.jncimonographs.a024234

62. Ball P, Emons G, Kayser H, Teichmann J. Metabolic clearance rates of catechol estrogens in rats. Endocrinology. 1983;113(5):1781–3. doi: 10.1210/endo-113-5-1781

63. Martucci C, Fishman J. Direction of Estradiol Metabolism as a Control of its Hormonal Action—Uterotrophic Activity of Estradiol Metabolites. Endocrinology. 1977;101(6):1709–15. doi: 10.1210/endo-101-6-1709

64. Martucci CP, Fishman J. Impact of continuously administered catechol estrogens on uterine growth and luteinizing hormone secretion. Endocrinology. 1979;105(6):1288–92. doi: 10.1210/endo-105-6-1288

65. Panek DU, Dixon WR. Effect of continuous intraventricular estrogen or catechol estrogen treatment on catecholamine turnover in various brain regions. J Pharmacol Exp Ther. 1986;236(3):646–52. doi: 10.1016/S0022-3565(25)38947-0

66. Parvizi N, Wuttke W. Catecholestrogens Affect Catecholamine Turnover Rates in the Anterior Part of the Mediobasal Hypothalamus and Medial Preoptic Area in the Male and Female Castrated Rat. Neuroendocrinology. 1983;36(1):21–6. doi: 10.1159/000123523

67. Al-Shami K, Awadi S, Khamees A, Alsheikh AM, Al-Sharif S, Ala’ Bereshy R, et al. Estrogens and the risk of breast cancer: A narrative review of literature. Heliyon. 2023;9(9):e20224. doi: 10.1016/j.heliyon.2023.e20224

68. Guillemette C, Bélanger A, Lépine J. Metabolic inactivation of estrogens in breast tissue by UDP-glucuronosyltransferase enzymes: an overview. Breast Cancer Res. 2004;6(6):246. doi: 10.1186/bcr936

69. Purohit A, Woo LWL, Potter BVL. Steroid sulfatase: A pivotal player in estrogen synthesis and metabolism. Mol Cell Endocrinol. 2011;340(2):154–60. doi: 10.1016/j.mce.2011.06.012

70. Tobiansky DJ, Korol AM, Ma C, Hamden JE, Jalabert C, Tomm RJ, et al. Testosterone and corticosterone in the mesocorticolimbic system of male rats: Effects of gonadectomy and caloric restriction. Endocrinology. 2018;159(1):450–64. doi: 10.1210/en.2017-00704

71. Wall EG, Desai R, Aung ZK, Yeo SH, Grattan DR, Handelsman DJ, et al. Unexpected Plasma Gonadal Steroid and Prolactin Levels Across the Mouse Estrous Cycle. Endocrinology. 2023;164(6):1–7. doi: 10.1210/endocr/bqad070

72. Fabregat-Safont D, Alechaga É, Haro N, Gomez-Gomez À, Velasco ER, Nabás JF, et al. Towards the non-invasive determination of estradiol levels: Development and validation of an LC-MS/MS assay for quantification of salivary estradiol at sub-pg/mL level. Anal Chim Acta. 2024;1331:343313. doi: 10.1016/j.aca.2024.343313

73. Balthazart J, Baillien M, Ball GF. Phosphorylation processes mediate rapid changes of brain aromatase activity. J Steroid Biochem Mol Biol. 2001;79:261–77. doi: 10.1016/S0960-0760(01)00143-1

74. Hayashi T, Harada N. Post-translational dual regulation of cytochrome P450 aromatase at the catalytic and protein levels by phosphorylation/dephosphorylation. FEBS J. 2014;281(21):4830–40. doi: 10.1111/febs.13021

75. Remage-Healey L, London SE, Schlinger BA. Birdsong and the neural production of steroids. J Chem Neuroanat. 2010;39(2):72–81. doi: 10.1016/j.jchemneu.2009.06.009

76. Micevych P, Dominguez R. Membrane estradiol signaling in the brain. Front Neuroendocrinol. 2009;30(3):315–27. doi: 10.1016/j.yfrne.2009.04.011

77. Lannigan DA. Estrogen receptor phosphorylation. Steroids. 2003;68:1–9. doi: 10.1016/S0039-128X(02)00110-1

78. de Bournonville MP, de Bournonville C, Vandries LM, Nys G, Fillet M, Ball GF, et al. Rapid changes in brain estrogen concentration during male sexual behavior are site and stimulus specific. Sci Rep. 2021;11(1):1–13. doi: 10.1038/s41598-021-99497-1

79. Wingfield JC, Wada M. Changes in plasma levels of testosterone during male-male interactions in the song sparrow, Melospiza melodia: time course and specificity of response. J Comp Physiol A. 1989;166(2):189–94. doi: 10.1007/BF00193463

